# High accuracy protein structures from minimal sparse paramagnetic solid-state NMR restraints

**DOI:** 10.1101/463158

**Authors:** Alberto Perez, Kari Gaalswyk, Christopher P. Jaroniec, Justin L. MacCallum

**Affiliations:** Department of Chemistry, University of Calgary, Calgary, Alberta, Canada; Department of Chemistry, University of Florida, Gainesville, Florida, USA; Department of Chemistry and Biochemistry, The Ohio State University, Columbus, Ohio, USA

**Keywords:** solid-state NMR, integrative structural biology, protein structure, computational modeling

## Abstract

There is a pressing need for new computational tools to integrate data from diverse experimental approaches in structural biology. We present a strategy that combines sparse paramagnetic solid-state NMR restraints with physics-based atomistic simulations. Our approach explicitly accounts for uncertainty in the interpretation of experimental data through the use of a semi-quantitative mapping between the data and the restraint energy that is calibrated by extensive simulations. We apply our approach to solid-state NMR data for the model protein GB1 labeled with Cu^2+^-EDTA at six different sites. We are able to determine the structure to *ca.* 1 Å accuracy within a single day of computation on a modest GPU cluster. We further show that in 4 of 6 cases, the data from only a single paramagnetic tag are sufficient to fold the protein to high accuracy.

---

Magic-angle spinning solid-state nuclear magnetic resonance (NMR) has emerged as a viable tool for analysis of the structure and dynamics of large biomacromolecular complexes and assemblies including membrane proteins, amyloids, viral capsids and chromatin^[1–5]^. In spite of the latest advances^[6,7]^, high-throughput protein structure elucidation by conventional solid-state NMR methods continues to be challenging due to inherent difficulties in obtaining critical non-local contacts. During the past decade, solid-state NMR measurements of nuclear paramagnetic relaxation enhancements (PREs) have been explored for structural studies of native metalloproteins^[8,9]^ and proteins modified with covalent paramagnetic tags, including nitroxide spin labels^[10]^ and metal chelates^[11–13]^. These paramagnetic methods are capable of providing information about electron-nucleus distances up to ~20 Å, providing valuable information for structure determination.

Notably, *de novo* solid-state NMR three-dimensional structure determination based on paramagnetic tagging in absence of internuclear distance restraints has been successfully demonstrated for the model 56-residue B1 immunoglobulin binding domain of protein G (GB1) in the microcrystalline phase^[11,14,15]^. Collectively, these studies highlight the considerable promise of such approaches. At the same time, they underscore the urgent need for more effective approaches toward generating accurate structural models based on paramagnetic restraints, as such approaches are expected to be key for future applications of this methodology to larger systems^[16,17]^.

Several challenges are present when modeling biomolecular structures based on PRE data for proteins modified with covalent tags. First, these measurements provide a limited number of independent experimental restraints—less than one per amino acid per paramagnetic mutant, as many of the measured distances are highly correlated. Second, it is desirable to limit the number of paramagnetic tagged protein variants in order to minimize the associated time, labor, and expense of sample preparation and data acquisition. Third, some of the distances derived from PRE measurements can be imprecise due to the presence of confounding effects, including intermolecular interactions, secondary metal binding sites, and diamagnetic contamination^[18,19]^. Finally, relaxation is affected by conformational heterogeneity, exacerbated by the inverse sixth power relationship between the PRE and the electron-nucleus distance^[19]^. While the last issue can in principle be addressed by ensemble refinement^[20]^, this comes at the expense of considerable complexity and computational cost.

Here, we employ a physics-based Bayesian approach called Modelling Employing Limited Data (MELD)^[21]^, which makes statistically consistent inferences about conformational ensembles by combining prior information (physical models of protein energetics and probe heterogeneity) with experimental data, enabling correct protein folding on timescales far faster than naïve molecular dynamics simulations would produce^[22]^. The approach is illustrated for the model protein GB1, for which extensive solid-state NMR PRE measurements are available for six EDTA-Cu^2+^ analogs^[11]^. We demonstrate that, for this protein, data from even a single paramagnetic label are often sufficient for accurate structure determination, compared to other approaches that require more data to achieve acceptable results.

We first developed a calibrated semi-quantitative mapping (Fig. 1) that turns ensemble averaged solid-state NMR PRE measurements into allowed distance ranges, which avoids the need for ensemble refinement. We performed molecular dynamics simulations (see Supporting Information) on a benchmark set of six small proteins, each labeled at ten different sites with a cysteine-EDTA-Cu^2+^ sidechain. From these simulations, we can predict the ^15^N longitudinal 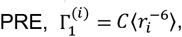 where the angle brackets denote the ensemble average, *r_i_* is the distance between the Cu^2+^ ion and the backbone amide nitrogen of residue *i*, and *C*=1.268×10^−54^ m^6^s^-1^ for the experimental conditions of this study. Based on experimental considerations, we divided the PREs into three strengths: *strong* 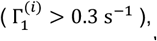 *medium* 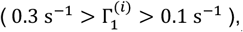 and *weak* 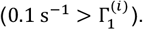 We then correlated the predicted 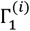 values to the corresponding distance between the Cα of the residue containing the EDTA-Cu^2+^ sidechain and the amide nitrogen of residue *i* (Fig. 1).

**Figure 1.**
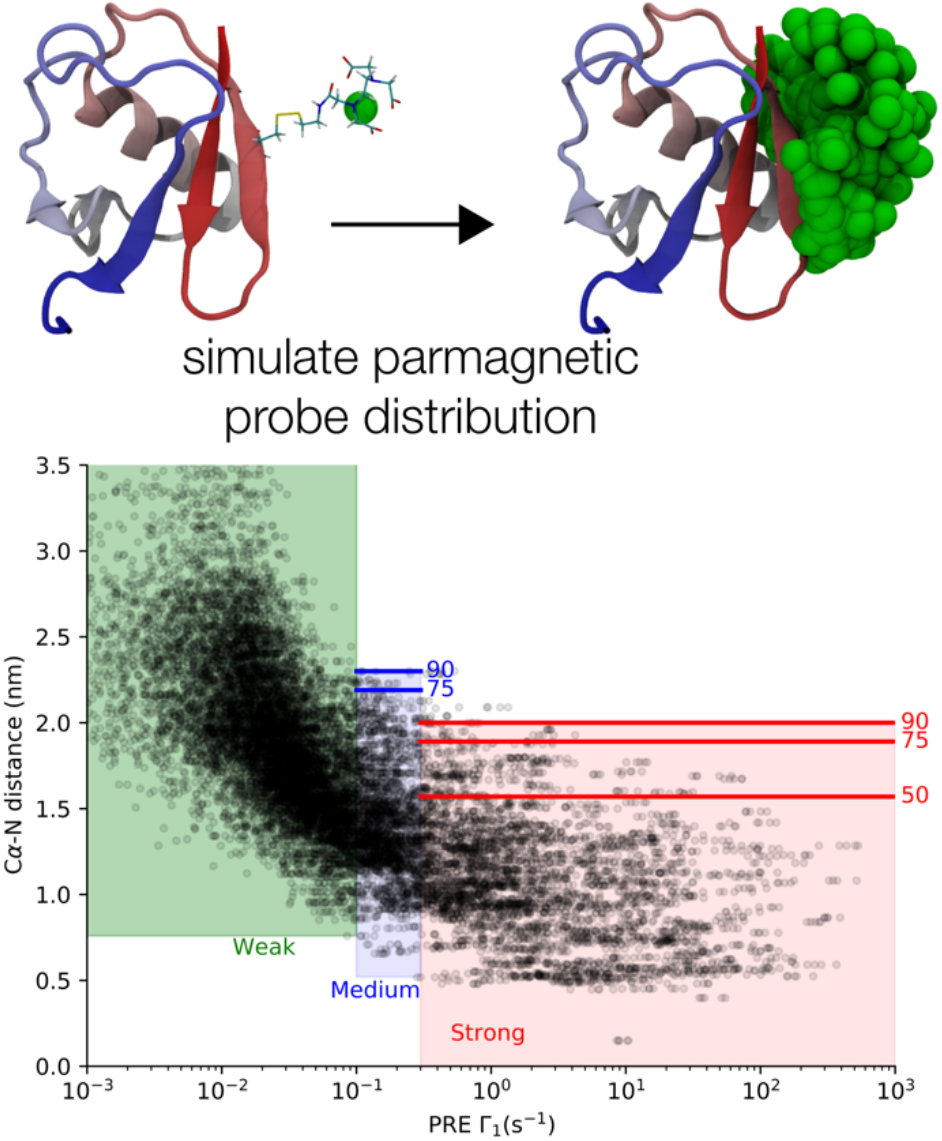
Semi-quantitative mapping from backbone amide 15N longitudinal PRE values (Γ_1_) to allowed distance ranges from ensemble average data. Upper: simulations were performed for a series of six small benchmark proteins each with ten different EDTA-Cu^2+^ tag locations. Lower: each point correlates the simulated Γ_1_ value to the corresponding C4-N distance. The computed Γ_1_ values were divided into strong, medium, and weak, and for each category, we identified allowable distance ranges encompassing 50, 75, or 90% of the Cα-N distances (see Supporting Information).

This map explicitly accounts for tag flexibility, leading to wide distance distributions (Fig. 1). Allowed distance ranges that encompass all observed distances would be overly broad, resulting in poor structures. However, making the allowed distance ranges too narrow would incorrectly penalize some structures, biasing the ensemble. To overcome this, we leverage the fact that MELD has the unique ability to handle unreliable restraints, where only a specified percentage of the restraints need be correct^[21,22]^. This allows us to specify that most distances must fall within a relatively narrow range, while a smaller number of distances can fall within a wider range (Table S1, see Supporting Information for details). The effects of the possible conformational heterogeneity are explicitly incorporated into the distance ranges, trading a loss of precision for the elimination of ensemble refinement.

We assessed the utility of this approach by structure determination of the protein GB1 (not used to derive the mapping), for which an extensive set (Fig. 2A) of solid-state NMR longitudinal backbone amide ^15^N PREs is available^[11]^. The measured 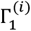 values were turned into allowed distance ranges and the protein secondary structure was predicted based on the GB1 backbone ^13^C and ^15^N chemical shifts using TALOS+^[23]^ (see Supporting Information).

**Figure 2.**
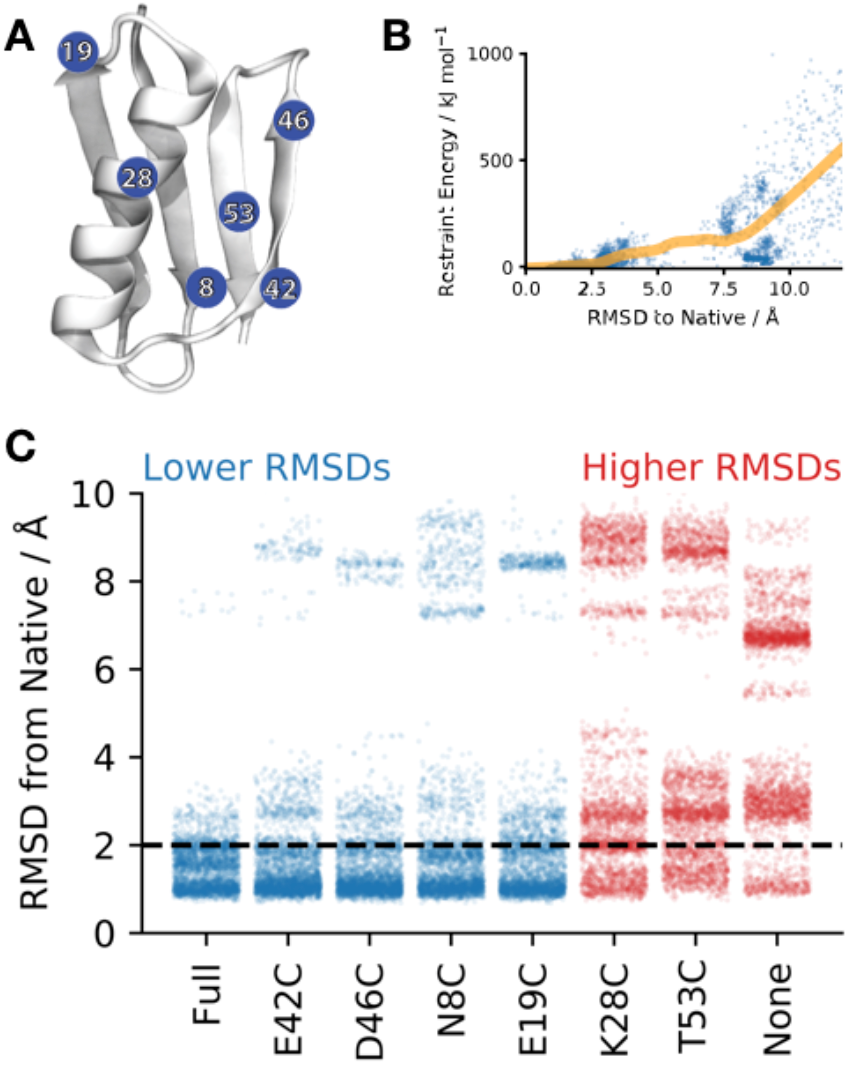
Accurate protein fold determination using MELD and sparse paramagnetic solid-state NMR restraints. (A) Location of the six EDTA-Cu^2+^ tag sites on the native structure of GB1. (B) Inclusion of the *Full* dataset results in an energy landscape that favours native-like structures. The restraint energy (from the *Full* dataset) and RMSD of all structures sampled in any of the simulations (blue dots) are shown, along with a LOESS regression (orange line). (C) RMSD distributions for all data sets explored in this work. Some data sets sample structures less than 2 Å backbone RMSD from the X-ray structure more than 70% of the time (blue) indicating that data for even a single paramagnetic mutant can be sufficient for accurate folding with MELD, whereas other datasets spend less than 30% of the time in near-native conformations (red).

To systematically assess how much PRE data is needed to generate accurate structures, we performed simulations with either all the available restraints (“*Full*”) or using PRE data for one mutant at a time, and compared these simulations to the case where no PRE data were used (“*None*”; still includes secondary structure restraints). Simulations were carried out in triplicate for 0.5 μs (< 24 hours each on 24 NVIDIA GTX 1080 Ti GPUs).

As expected, the *Full* dataset produces the narrowest ensembles (Fig. 2C), with over 70% of the sampled structures within 2 Å backbone RMSD of the crystallographic structure (PDB ID: 2gb1). The introduction of PRE data has a strong funneling effect (Fig. 2B), where structures disagreeing with the data have large energetic penalties. Clustering of the *Full* dataset results in a dominant cluster with protein conformations that have ~1 Å RMSD from the GB1 crystal structure. Remarkably, the resulting protein structures display excellent sidechain packing (Fig. 3), even though the experimental data report only on distances between the paramagnetic tag and backbone amide nitrogen atoms. The restraint energy is flat within about 2.5 Å of the crystal structure (Fig. 2B), so the accuracy of the structural model to within 1 Å and the correct packing of the core sidechains can be attributed to the accuracy of the physical model.

**Figure 3.**
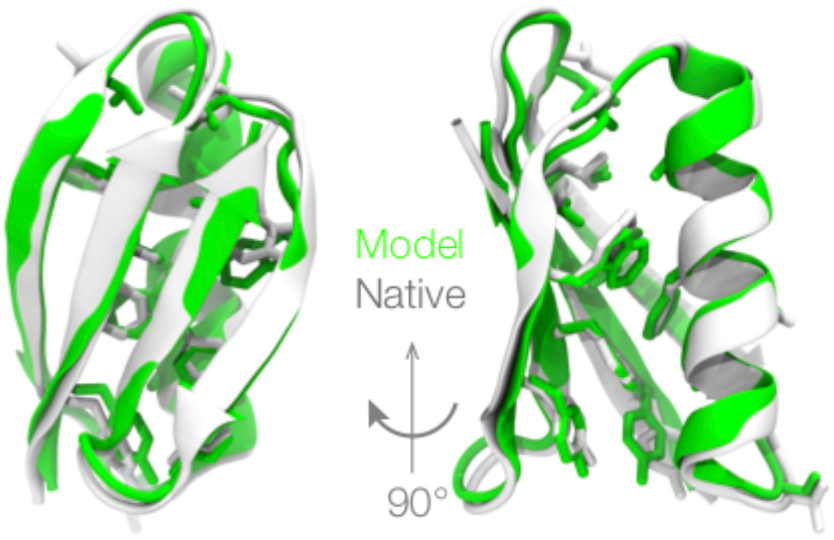
Most conformations exhibit accurate sidechain packing, even though the experimental data report only on backbone distances. A representative structure from the dominant cluster of the *Full* dataset (green) is shown superimposed on the GB1 crystal structure (white), with the core sidechains shown as sticks. The backbone RMSD is ~0.9 Å.

The individual datasets for most EDTA-Cu^2+^ locations (*E42C*, *D46C*, *N8C*, *E19C*) also lead to tight ensembles around the native state with more than 70% of the sampled conformations within 2 Å backbone RMSD from the X-ray structure. In contrast, *K28C* and *T53C* produce ensembles that are only somewhat improved over *None*. For comparison, standard unbiased MD simulations using a closely-related force-field and solvation model did not sample any near-native structures within 50 μs, which is 100 times longer than the current simulations^[24]^.

The *E42C* dataset is representative of the four datasets that have strongly funneled restraint energy landscapes and produce accurate models (Fig. 4). These datasets feature strong, non-local PREs that restrain residues that are far apart in the sequence, greatly reducing the available conformational entropy^[25]^. Although the restraint energy landscape is flat within ~5 Å of the crystal structure, nearly all models are within 2 Å of the crystal structure, which we attribute to the quality of the physical modeling and sampling approach.

**Figure 4.**
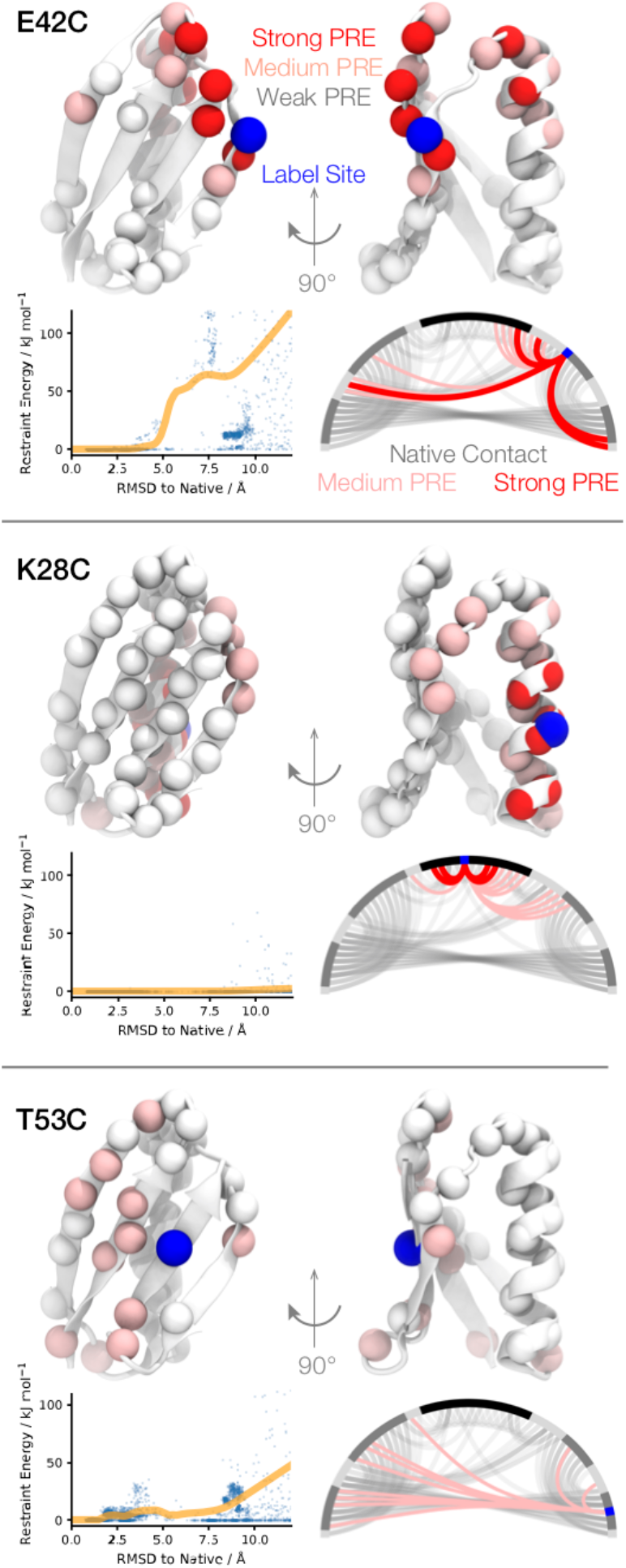
Probe locations that give strong, non-local PREs are more informative and give more funnelled energy landscapes. Top row: each panel shows the strong (red), medium (pink), or weak (white) PREs mapped onto the structure. Lower left: the restraint energy for the current dataset and RMSD are shown for all structures sampled in any of the simulations in this study (blue dots), along with a LOESS regression (orange line). Lower right: The native contacts (C*α*-C*α* distance < 10 Å, grey), and strong (red) and medium (pink) PRE restraints mapped onto the protein sequence. Grey segments along the perimeter denote secondary structural elements.

The *K28C* dataset produces a uniformly flat restraint energy landscape (Fig. 4). This dataset provides scant non-local information regarding the proximity of different secondary structure elements. Instead, it yields primarily redundant information about contacts within the *α*-helix that provide little additional information beyond the TALOS+ secondary structure predictions.

We anticipated that *T53C* would be highly informative, as residues 5–7 are directly adjacent to residue 53, and we thus expected *strong* PREs that would serve to restrain the terminal β- strands. However, the measured PREs were somewhat weaker (~0.1–0.2 s^-1^) than expected, falling into the *medium* category, possibly due to diamagnetic contamination of the sample, where the EDTA tags are not completely saturated with Cu^2+^, although we have not established this definitively. These weaker than expected PREs lead to an energy landscape that disfavors some near-native conformations, contributing to the broad ensemble observed. This result highlights the need for caution in over-interpreting PRE data, given the complications from intermolecular interactions, secondary metal binding sites, conformational heterogeneity, and diamagnetic contamination, and underscores the need for modeling approaches, such as MELD^[21,22]^ and analogous methods^[14, 26–28]^, which are able to deal with sparse and imprecise experimental data.

In the absence of a reference protein structure would it be possible to predict which calculations are accurate and which are not? Observation of the same dominant conformation from multiple datasets, as observed here, provides a strong indication of reliability. We also observed that simulations with tight ensembles were more likely to be accurate (Figure S1).

Previous studies have assessed the utility of solid-state NMR PRE restraints for structure determination on the same system (GB1), which enables direct comparison. Sengupta et al.^[11]^ used Xplor-NIH^[29]^ in combination with the same dataset as the present study. Using all six EDTA-Cu^2+^ sites, totaling ~230 measurements, they were able to produce a model with a backbone RMSD of 1.8 Å relative to the crystal structure. Even when using the complete set of PREs, the convergence properties of the Xplor-NIH based approach were relatively poor, requiring the calculation of many hundreds of protein structures within a two-stage refinement procedure to generate the final ensemble of low energy models. Combining our semi-quantitative MELD approach with the same dataset results in a much tighter structural ensemble (Fig. 2, “*Full”*), and a representative model that is 0.9 Å from native with excellent sidechain packing (Fig. 3). Furthermore, for GB1, MELD is able to produce similarly accurate models using data from only a single EDTA-Cu^2+^ mutant corresponding to a total of ~30-40 PRE restraints (Fig. 2). Following a closely related experimental approach, Tamaki et al.^[15]^ used a CS-Rosetta^[30]^ based protocol to fold GB1 to within 1.5 Å of the native structure using three paramagnetic mutants. The use of PRE data from single EDTA-Mn^2+^ analogs resulted in structural models with RMSDs in the 4.5–8 Å regime, which is substantially worse than MELD’s performance.

In summary, MELD can be used to generate highly accurate protein structural models based on sparse solid-state NMR PRE restraints, while successfully overcoming several challenges related to heterogeneity. MELD is able to produce accurate results using only a small fraction of the experimental data, which translates to substantial potential time and cost savings, promising to considerably increase the throughput of solid-state NMR structural studies of proteins. These results further highlight the increasing power and utility of integrative methods in biomacromolecular structure determination, even in the limit of small amounts of information. We expect that such approaches will continue to gain popularity, particularly in cases where sparse structural data can be obtained for systems that are not amenable to analysis by the traditional structural biology techniques.

## Experimental Section

Residue-specific solid-state NMR longitudinal ^15^N PRE data were recorded for six isostructural cysteine-EDTA-Cu^2+^ mutants of GB1 as described previously^[11]^. Simulations were carried out using the MELD^[21]^ software package. Full computational details can be found in the Supporting Information.

## Acknowledgements

JLM is a Canada Research Chair. This work was supported by the National Science Foundation (grant MCB-1715174 to CPJ), the Camille & Henry Dreyfus Foundation (CPJ), the Natural Sciences and Engineering Research Council of Canada (JLM), and Compute Canada (JLM).

## 1 Development of Semi-Quantitative Map

### 1.1 Calibration Simulations

To derive the semi-quantitative mapping as described in the main text, benchmark simulations of six small proteins (1cqg, 1d3z, 1erc, 1nkl, 1ubq, 6lyz) ^[1–6]^ were carried out. Ten simulations of each protein were performed, each with a different surface-exposed residue mutated to a Cys-EDTA-metal sidechain. The Cys-EDTA sites were chosen to encompass a mixture of helices, strands, and loops, all at surface exposed positions that would not be expected to cause major perturbation to the structure. Simulations were 100 ns long with a 2 fs timestep. All bonds involving hydrogen were constrained to their ideal lengths. The temperature was maintained at 300 K. Proteins were modelled using the Amber ff14SB force field^[7]^; the Cys-EDTA sidechain was modelled using the General Amber Force Field^[8]^; and the solvent was modelled using the GBneck2 implicit model^[9]^.

A few simulations displayed some instability, where the conformation occasionally drifted away from the crystal structure, likely due to limitations of the combination of force field and solvation model^[10]^. To mitigate this, any conformations that were more than 3 Å from the crystal structure were excluded from the analysis.

### 1.2 Prediction of Relaxation Enhancement

Metal–nitrogen distance distributions were converted into simulated longitudinal ^15^N PRE values (Γ_1_) through the Solomon-Bloembergen equations:

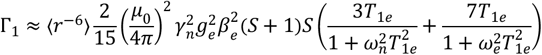

where *r* is the electron-nucleus distance, *μ*_0_ is the permeativity of free space, *γ_n_* is the nuclear gyromagnetic ratio, 34is the electronic g-factor, *β_e_* is the Bohr magneton, *S* is the electron spin quantum number of the paramagnetic ion, *ω_n_* is the Larmor frequency of the nucleus, *ω_e_* is the Larmor frequency of the electron, and *T*_1e_ is the effective longitudinal relaxation time constant.

This can be simplified to

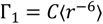

where *C*=1.268×10^−54^ m^6^s^-1^ under the experimental conditions here (external magnetic field of 11.7 T, effective electron longitudinal relaxation time constant for Cu+ of 2.5 ns).

Statistical analysis of the experimental Γ_B_ measurements showed noise levels of approximately 10 percent of the measured Γ_B_ values due to instrumental noise and data processing artifacts. To simulate this, we generated 10 samples of each Γ_B_ value by adding zero-mean Gaussian noise with a coefficient of variation of 0.1.

We then correlated the predicted 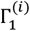 values to the corresponding distance between the CD of the residue containing the EDTA-Cu^2+^ sidechain and the amide nitrogen of residue *i* (Fig. 1).

### 1.2 Extraction of Distance Ranges

Based on experimental considerations, we divided the PREs into three strengths: *Strong* 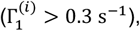 *Medium* 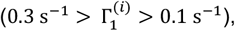 and *Weak* 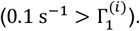 We extracted unreliable distance ranges from the benchmark simulations with different degrees of reliability. For each PRE strength and each EDTA-Cu^2+^ tag/protein combination, we derived a lower bound that encompasses all observed CD-N distances. We derived a set of upper bounds that encompass various percentages (*strong*: 50, 75, and 90, *medium*: 75 and 90, *weak*: 100) of the observed distances. We then took the broadest distance ranges for each strength across all protein and tag location combinations, establishing the worst-case distance bounds for each PRE strength and percentage (Table S1). These are the tightest distance ranges that have zero energy for every tag location in all of our benchmark simulations.

**Table S1.** Distance bounds derived from benchmark simulations of 6 small proteins at 10 different tag locations each.

| Strength<sup>[a]</sup> | Percentage Reliability<sup>[b]</sup> | Lower Bound (Å) | Upper Bound (Å)<sup>[c]</sup> |
| --- | --- | --- | --- |
| Strong | 50 | 0.0 | 15.7 |
|  | 75 | 0.0 | 18.9 |
|  | 90 | 0.0 | 20.0 |
| Medium | 75 | 5.2 | 21.5 |
|  | 90 | 5.2 | 23.0 |
| Weak | 100 | 7.6 | ∞ |

[a] *Strong*: ( $\Gamma_1 > 0.3 \text{ s}^{-1}$ ), *Medium*: ( $0.3 \text{ s}^{-1} > \Gamma_1 > 0.1 \text{ s}^{-1}$ ), *Weak*: ( $0.1 \text{ s}^{-1} > \Gamma_1$ ). [b] Fraction of restraints that are enforced during MELD simulation.

## 2 Structure Determination Calculations

### 2.1 Treatment of Experimental PRE Data

The relaxation enhancement was quantified by fitting Γ_1_ to experimental relaxation measurements as described previously[11]. Allowed distance ranges between the CD of the labelled residue and the measured amide nitrogen were calculated based on the magnitude of Γ_1_ (Table S1).

For each spin label location:

- All of the residue pairs corresponding to *Strong* restraints were placed into a “collection”.
- 50, 75, and 90% of the distances were restrained to fall within the ranges given in Table S1
- The same process was repeated for *Medium* and *Weak* restraints

We note that the collections for each paramagnetic probe are independent, so that the specified percentage of distance bounds must be satisfied for each probe individually.

### 2.2 Treatment of Experimental Backbone Chemical Shift Data

Secondary structure information was incorporated through TALOS+ derived restraints based on backbone ^13^C and ^15^N chemical shifts for GB1^[12]^ (BMRB entry 15156). The protocol for secondary structure restraints in MELD was as descried previously^[13]^.

### 2.3 Setup of MELD Simulations

The MELD algorithm^[13]^, based upon the open-source GPU-accelerated OpenMM simulation engine^[14]^, was used to generate structural models from the collected NMR data. All simulations were 500 ns long and were repeated in triplicate with different random seeds. Proteins were modeled using the ff14SB force field^[7]^ plus AMAP torsion potential correction^[15]^ with the GBneck2 implicit solvation model^[15]^. A combined Hamiltonian and temperature replica exchange approach was used^[13]^ with 24 replicas. The force constants for PRE restraints were “turned on” from zero to 250 kJ mol^-1^ nm^-2^ from replica 24 to 12, with *Strong* PREs turned on from 24–17 and *Medium and Weak* PREs turned on from 17–12. The temperature was held constant at 550 K in this region. The force constants remained at full strength in replicas 1–12, whereas the temperature was geometrically decreased from 550 K at replica 12 to 300 K at replica 1. Replica exchanges were attempted every 50 ps.

## 3 Analysis

### 3.1 More Accurate Calculations Display Tighter Ensembles

Our physics-based approach was able to produce highly accurate structural models even when using PRE data from only a single mutant for four of the six EDTA-Cu^2+^ mutants of GB1 (except K28C and T53C). This finding raises a question of whether in the absence of a reference protein structure it is possible to predict which calculations are accurate and which are not? There are a number of possible strategies to address this.

As described in the main text, one possibility is to assess the consistency of models produced using different datasets. In this work, the dominant conformers produced in the *Full*, *N8C*, *E19C*, *E42C*, and *D46C* datasets are highly similar. These conformations are even present in the other datasets, although far less frequently (Fig. 2). This is a form of cross-validation, and the observation of the same dominant conformation across a range of datasets lends confidence to the accuracy of the models.

Another possibility is to examine the “width” of the ensembles (Fig. S1), quantified as the mean pairwise RMSD between ensemble members, compared to the accuracy of the ensemble (quantified by the average RMSD to native). There is a clear correlation between ensemble width and model accuracy. When the ensemble is broad, it means that MELD is uncertain about what the structure is. Assuming the data is for a well-structured protein, it is unlikely that the predictions are accurate under these circumstances. On the other hand, when the ensemble is narrow, it means that MELD is quite certain about what the structure is. While this does not guarantee that the structure is accurate, it at least provides some indication of MELD’s confidence in the prediction. These observations are consistent with previous experiences^[13,16,17]^, where we generally find a correlation between the “tightness” and accuracy of the ensemble.

**Figure S1:**
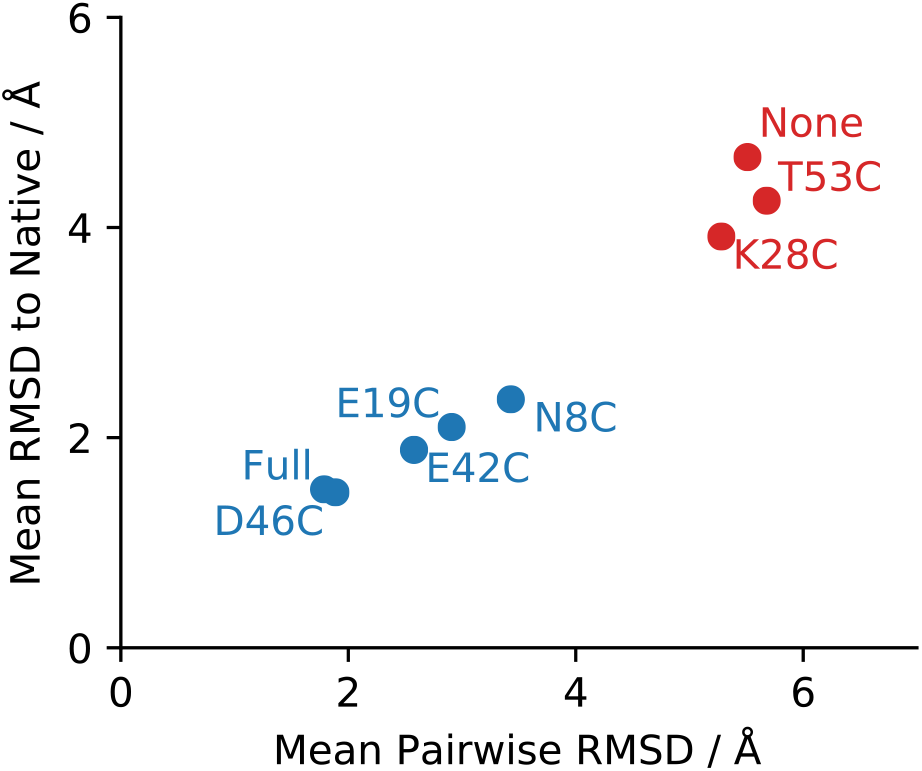
Mean pairwise RMSD within the simulated ensemble predicts model accuracy. Data set are colored depending on if they spend more (>70%, blue) or less (<30%; red) time in near-native (<2 Å backbone RMSD from X-ray) conformations. Data sets with broad ensembles (further to the right) contain lower quality models with higher RMSD to the native structure.

